# The ion channel Trpc6a regulates the cardiomyocyte regenerative response to mechanical stretch

**DOI:** 10.1101/2023.03.14.532536

**Authors:** Laura Rolland, Adèle Faucherre, Jourdano Mancilla Abaroa, Aurélien Drouard, Chris Jopling

**Author notes:** Correspondance: Chris Jopling.

## Abstract

Myocardial damage caused for example by cardiac ischemia leads to ventricular volume overload resulting in increased stretch of the remaining myocardium. In adult mammals, these changes trigger an adaptive cardiomyocyte hypertrophic response which, if the damage is extensive, will ultimately lead to pathological hypertrophy and heart failure. Conversely, in response to extensive myocardial damage, cardiomyocytes in the adult zebrafish heart and neonatal mice proliferate and completely regenerate the damaged myocardium. We therefore hypothesized that in adult zebrafish, changes in mechanical loading due to myocardial damage may act as a trigger to induce cardiac regeneration. Based, on this notion we sought to identify mechanosensors which could be involved in detecting changes in mechanical loading and triggering regeneration. Here we show using a combination of knockout animals, RNAseq and *in vitro* assays that the mechanosensitive ion channel Trpc6a is required by cardiomyocytes for successful cardiac regeneration in adult zebrafish. Furthermore, using a cyclic cell stretch assay, we have determined that Trpc6a induces the expression of components of the AP1 transcription complex in response to mechanical stretch. Our data highlights how changes in mechanical forces due to myocardial damage can be detected by mechanosensors which in turn can trigger cardiac regeneration.

## Introduction

Following a myocardial infarction, the loss of cardiac tissue results in significant changes in the mechanical loads exerted on the heart. As dynamic/elastic myocardium is replaced by relatively stiff non-contractile scar tissue, the resulting elevated preload and changes in tissue composition increases the amount of stretch exerted on the remaining myocardium, triggering an increased force of contraction and concomitant cardiomyocyte hypertrophy (Frank-Starling law)^1^. While this adaptive mechanism initially compensates for the increased mechanical load, prolonged stress will become maladaptive and, without medical intervention, will ultimately result in pathological hypertrophy and heart failure. In contrast to adult mammals, neonatal mice and adult zebrafish can fully regenerate their hearts after cardiac injury^2,3^. Rather than undergoing compensatory hypertrophy, neonatal mouse and adult zebrafish cardiomyocytes proliferate in response to the loss of myocardium. This suggests that the changes in mechanical loading which occur after cardiac injury in neonates/zebrafish may act as a trigger to induce cardiomyocyte proliferation and ultimately regeneration. Although decades of research have been devoted to understanding the effects of mechanical stretch on non-proliferative adult cardiomyocytes there is very little information regarding the effects on cardiomyocytes which are capable of proliferation. However, *in vitro* evidence studying the effects of mechanical loading on embryonic mouse cardiomyocytes indicates an increase in the proliferation index following 24 hours of cyclic stretch^4^. Likewise, transcriptomic analysis of neonatal rat ventricular cardiomyocytes subjected to cyclic stretch *in vitro* showed a significant upregulation of genes associated with cell proliferation^5^. To understand whether increased mechanical stretch can induce cardiomyocyte proliferation *in vivo*, in models which are capable of cardiac regeneration, it is important to assess whether changes in mechanical force occur following cardiac injury. Analysis of adult zebrafish indicates that following cardiac injury, the initial response is similar to that observed in adult humans resulting in a stiffer myocardium due to the extensive fibrosis at the site of injury^6^. Furthermore, the resulting volume overload initially induces an elongation of cardiomyocyte sarcomere length which reverts following cardiac regeneration. This indicates that following cardiac injury in adult zebrafish, the heart is subjected to significant changes in mechanical loading. Whether these forces can induce a proliferative response is at present unclear^6^. However, other situations that can also increase cardiomyocyte stretch do support this hypothesis. In humans vigorous exercise results in an elevated preload resulting in increased cardiomyocyte stretch. These conditions trigger an adaptive hypertrophic response resulting in an essentially larger more powerful heart^1^. On the other hand, in adult zebrafish, exercise can induce cardiomyocytes to proliferate rather than undergo hypertrophy^7^. This indicates that increased mechanical load can act as a stimulus to induce cardiomyocyte proliferation. But care must be taken when extrapolating exercise induced cardiomyocyte hypertrophy vs damage induced hypertrophy as it appears that different mechanisms are at play depending on the conditions^8^. Therefore, understanding how increased stretch can influence cardiomyocyte proliferation, particularly *in vivo*, may provide invaluable information on how this phenomenon could be harnessed to induce a regenerative response. Increased myocardial stretch can be sensed by a wide variety of mechanosensory mechanisms present in cardiomyocytes such as cell surface receptors, sarcomeric components, intercalated discs and stretch activated ion channels^1^. The transient receptor potential (Trp) channels are a family of non-selective cation channels which can be regulated by a variety of stimuli including mechanical stretch. Of these, *TRPC3* and *TRPC6* are highly expressed in the heart and have been directly linked to pathological cardiac hypertrophy in response to chronic overload. In particular, stretch elevates intracellular Ca^2+^ which activates the CALCINEURIN/NUCLEAR FACTOR OF ACTIVATED T CELLS (NFAT) pathway triggering pathological hypertrophy and remodelling^9,10^. Both *TRPC3* and *TRPC6* have been shown to be responsible for the stretch induced increase in Ca^2+9^. Indeed, targeting both of these ion channels can inhibit pathological cardiomyocyte hypertrophy^11^. Because of the role TRPC6 plays in regulating the cardiomyocyte hypertrophic response to increased stretch in mammals we surmised that Trpc6 may also regulate cardiomyocyte proliferation in adult zebrafish regenerating hearts in response to the chronic mechanical overload associated with cardiac injury.

In this study we examined how the loss of *trpc6a* could affect cardiac regeneration in zebrafish. Similar to reports in mammals, we found that *trpc6a* knockout (KO) did not result in any observable cardiac developmental defects. However, adult *trpc6a* KO zebrafish failed to regenerate their hearts following apical resection due to a reduction in cardiomyocyte proliferation. Transcriptomic analysis indicated that following cardiac injury, *trpc6a* KO zebrafish did not upregulate the expression of genes required for this process. In particular, this included orthologous components of the ACTIVATOR PROTEIN 1 (AP1) transcription factor complex which is critical for cardiac regeneration in adult zebrafish^12^. Furthermore, we demonstrate that the stretch induced expression of AP1 components is markedly reduced in *trpc6a* KO zebrafish. Taken together these findings indicate that Trpc6a positively regulates the expression of pro-regenerative genes in response stretch and that this process is required for successful cardiac regeneration.

## Results

### Loss of *trpc6a* does not affect cardiac development

To understand the role Trpc6a plays during cardiac development and regeneration, we utilised a KO zebrafish line which harbours a single base pair substitution G637T resulting in a premature stop codon in exon2 of *trpc6a* (Fig.1.A). To confirm that this mutation results in a loss of Trpc6a protein, we performed immunohistochemistry (IHC) using a Trpc6 antibody. In this manner we could detect Trpc6 in the myocardium of adult *trpc6a*^*+/+*^ zebrafish hearts (Fig.1.B). In comparison, Trpc6 was absent in adult *trpc6a*^*-/-*^ zebrafish hearts indicating that no functional Trpc6a protein was present in the KO line (Fig.1.C). In parallel, we also assessed sarcomere structure in *trpc6a*^*-/-*^ using a Tropomyosin (trpm) antibody. We could not detect any observable differences in tropomyosin labelling between *trpc6a*^*-/-*^ and their *trpc6a*^*+/+*^ siblings (Fig.1.D,E). We next determined whether loss of Trpc6a affects cardiac development in zebrafish larvae as this could potentially impact processes occurring in adulthood, such as cardiac regeneration. Because of the role TRPC6 plays in hypertrophy in mammals we first measured the ventricular wall of 3 days post fertilisation (dpf) larvae (n=10/group) (Fig.1.F,G). In this manner we could not detect any significant differences between *trpc6a*^*-/-*^ and *trpc6a*^*+/+*^ larvae (Fig.1.H). Mutations in *TRPC6* have also been associated with cardiac arrhythmias which could, if present, disrupt heart regeneration at later stages. Therefore, we analysed a variety of cardiac physiological parameters in both *trpc6a*^*-/-*^ and *trpc6a*^*+/+*^ larvae. Measurements of ventricular and atrial heart rates indicated there was no significant difference between *trpc6a*^*-/-*^ and *trpc6a*^*+/+*^ larvae (n=10/group) (Fig.1.I). Lastly, we measured the blood flow rate and calculated the cardiac output in both *trpc6a*^*-/-*^ and *trpc6a*^*+/+*^ larvae. Our data indicates that there are no significant differences in these parameters between *trpc6a*^*-/-*^ and *trpc6a*^*+/+*^ larvae indicating that loss of Trpc6a does not appear to affect overall cardiac performance (n=10/group) (Fig.1.J,K). These data indicate that heart development and cardiac performance are not significantly affected in *trpc6a*^*-/-*^ zebrafish larvae.

**Figure 1.**
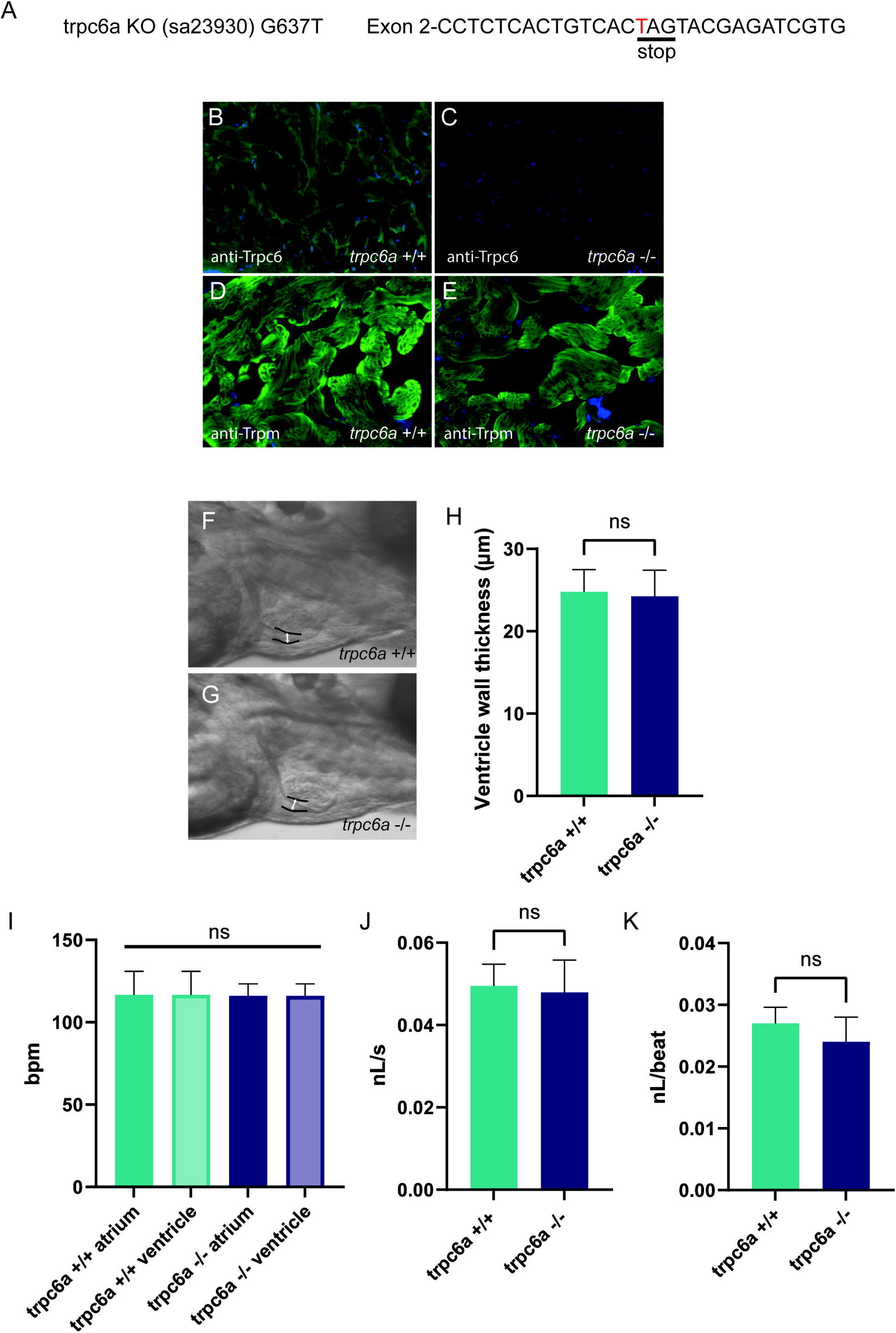
Loss of *trpc6a* does not affect cardiac development. **A**. Description of the point mutation carried by the sa23930 zebrafish transgenic line. The G637T mutation causes a premature stop codon in the Exon 2 of the *trpc6a*. **B-C**. IHC images from adult heart sections showing the presence of Trpc6 in the myocardium of *trpc6a*^+/+^ but absent from the myocardium of *trpc6a*^-/-^ zebrafish. Trpc6: green, DAPI: blue. **D-E**. IHC images from adult heart sections showing the organization of tropomyosin (Trpm) in the myocardium of *trpc6a*^*+/+*^ and in *trpc6a*^*-/-*^ zebrafish. Trpm: green, DAPI: blue. **F-G**. Representative morphology of the ventricular wall of 3dpf larvae from control (*trpc6a*^+/+^, **F**) and trpc6a KO (*trpc6a*^*-/-*^, **G**) groups. **H**. Ventricular wall thickness measurements of 3dpf larvae during diastole. t-test was used for statistical analysis. **I**. Atrial and ventricular contraction rate (in bpm) of *trpc6a*^+/+^ and *trpc6a*^-/-^ 3dpf larvae. 1-way ANOVA was used for statistical analysis. **J**. Blood flow velocity (in nL/s) measured in the caudal vein at 3dpf. t-test was used for statistical analysis. **K**. Cardiac output (in nL/beat) of *trpc6a*^+/+^ and *trpc6a*^-/-^ 3dpf larvae. t-test was used for statistical analysis. **F-K**. Data obtained on n=10 larvae per group.

### Trpc6a is required for cardiac regeneration

Because of the role TRPC6 plays in sensing changes in mechanical load after cardiac injury in mammals, we assessed whether the loss of Trpc6a affected cardiac regeneration in adult zebrafish. To achieve this, we performed apical resection of adult *trpc6a*^*-/-*^ and *trpc6a*^*+/+*^ zebrafish. At 30 days post amputation (dpa), histological staining using acid fuchsin orange G (AFOG) indicated that the loss of Trpc6a inhibited cardiac regeneration resulting in the presence of a significant fibrin/collagen scar (n=5/group) (Fig.2.A-C). Previous research indicates that TRPC6 can also play a role in angiogenesis^13^. During cardiac regeneration in adult zebrafish, revascularization of the wound region is a critical early event which could be affected by the loss of Trpc6a. To address this possibility, we analysed wound revascularization at 7dpa. In this manner we could detect numerous vessels forming a vascular plexus within the wound region of both *trpc6a*^*-/-*^ and *trpc6a*^*+/+*^ zebrafish hearts indicating that this process appears largely unaffected and is unlikely the cause of the defective regeneration we observed in *trpc6a*^*-/-*^ hearts (n=5/group)(Fig.2.D-G). We next sought to determine whether cardiomyocyte proliferation had been affected in *trpc6a*^*-/-*^ zebrafish. To meet this end, we preformed EdU labelling of resected *trpc6a*^*-/-*^ and *trpc6a*^*+/+*^ zebrafish hearts at 14dpa. Our analysis indicates that there is a significant reduction in the number of EdU labelled cardiomyocytes in the *trpc6a*^*-/-*^ hearts compared to their *trpc6a*^*+/+*^ siblings (n=3/group)(Fig.2.H-J). Taken together this data indicates that the loss of *trpc6a* disrupts cardiac regeneration due to a significant reduction in cardiomyocyte proliferation.

**Figure 2.**
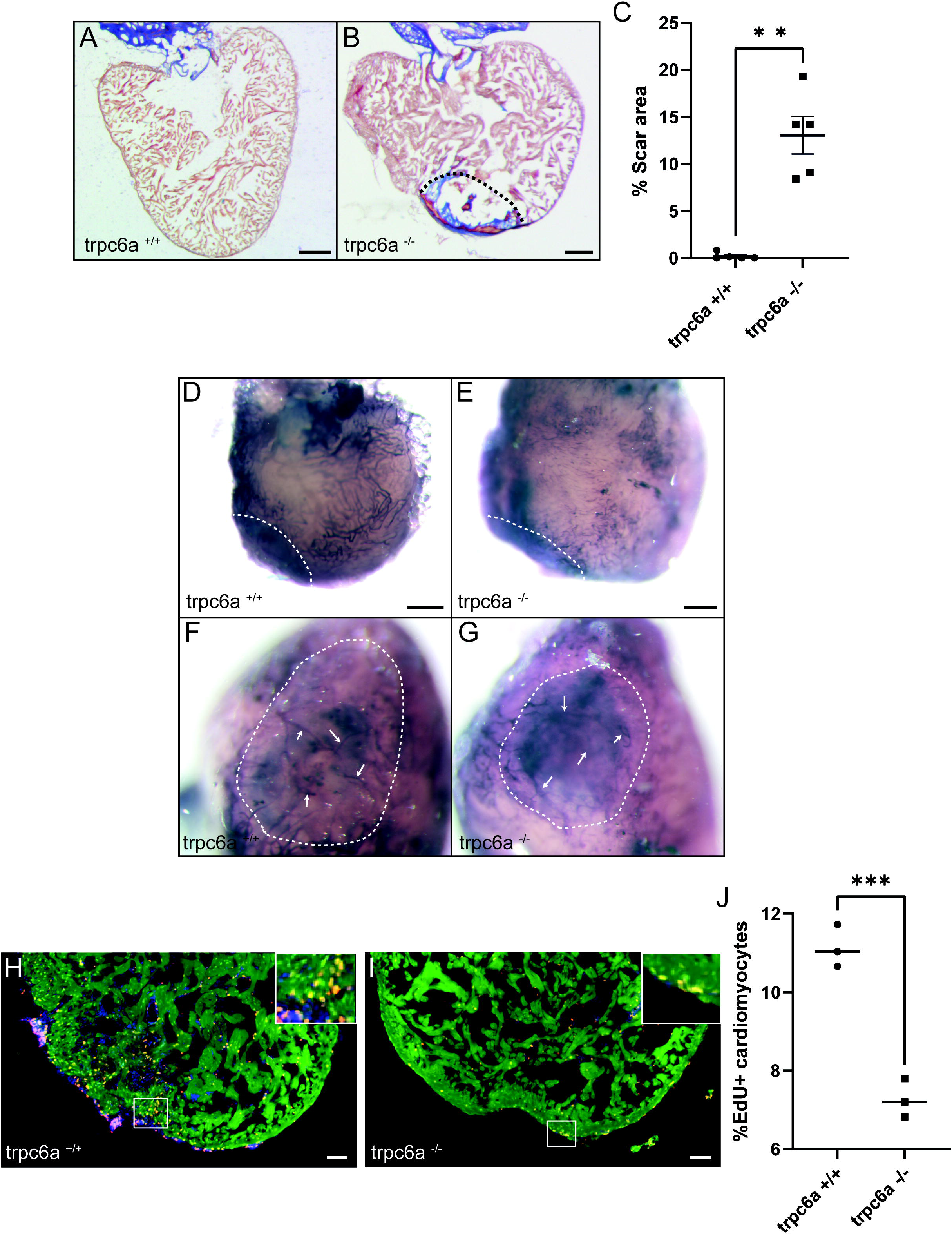
Trpc6a is required for cardiac regeneration. **A-C**. AFOG staining images and quantification of the scar area at 30dpa. Representative image of AFOG staining obtained for *trpc6a*^*+/+*^ (**A**) and *trpc6a*^*-/-*^ (**B**). Scale bars: 200μm. The dashed line outlines the scar region. The quantification of the scar area (n=5/group) (**C**). t-test was used for statistical analysis. **: p value < 0.01. **D-G**. Representative images of alcaline phosphatase staining showing the vasculature on 7dpa whole mount hearts. Low (**D-E**) and high (**F-G**) magnification of the vascular plexus present in the wound region of *trpc6a*^*+/+*^ (**D, F**) and *trpc6a*^*-/-*^ (**E, G**) zebrafish heart. Scale bars: 200μm. **H-J**. Cardiomyocyte proliferation measured at 14dpa. Representative IHC images showing Mef2c (green), EdU (red) and DAPI (blue) for *trpc6a*^*+/+*^ (**H**) and *trpc6a*^*-/-*^ (**I**). The white box depicts a higher magnification image in the upper right corner. Scale bars: 100μm. **J**. Quantification of EdU+ cardiomyocytes (n=3/group). t-test was used for statistical analysis. ***: p value < 0.001.

### Loss of Trpc6a results in misregulated of gene expression during regeneration

To determine what effect the loss of Trpc6a had on the cardiac transcriptome during heart regeneration, we performed bulk RNA sequencing of sham operated and 7dpa resected *trpc6a*^*-/-*^ and *trpc6a*^*+/+*^ hearts (n=5/group). Analysis of sham vs 7dpa samples indicates that 480 genes are significantly upregulated in 7dpa *trpc6a*^*+/+*^ hearts compared with 293 in 7dpa *trpc6a*^*-/-*^ (Fig.3.A-C). Of these, 135 genes are upregulated in both *trpc6a*^*-/-*^ and *trpc6a*^*+/+*^ 7 dpa hearts while 345 genes are specifically upregulated in *trpc6a*^*+/+*^ hearts and 158 genes are specific to *trpc6a*^*-/-*^ hearts (Fig.3.C). Analysis of significantly downregulated genes indicates that 8 are common to both *trpc6a*^*-/-*^ and *trpc6a*^*+/+*^ 7 dpa hearts while 48 are specifically downregulated in *trpc6a*^*+/+*^ hearts compared with 68 downregulated genes specific to *trpc6a*^*-/-*^ hearts (Fig.3.A-C). Lastly, 1 gene was significantly upregulated in *trpc6a*^*-/-*^ hearts and downregulated in *trpc6a*^*+/+*^ hearts while 2 genes were significantly upregulated in *trpc6a*^*+/+*^ hearts and downregulated in *trpc6a*^*-/-*^ hearts (Fig.3.C). We next focused on the 345 genes which were specifically upregulated in *trpc6a*^*+/+*^ hearts as these likely included genes which are induced following Trpc6a activation. In this manner we identified a number of transcription factors which were significantly upregulated in the *trpc6a*^*+/+*^ hearts but not in *trpc6a*^*-/-*^ hearts (Fig.3.D). Of particular interest were a number of components of the AP1 transcription factor complex (*fosl1a, june* and *fosab*), a critical regulator of cardiac regeneration in adult zebrafish^12^.

**Figure 3.**
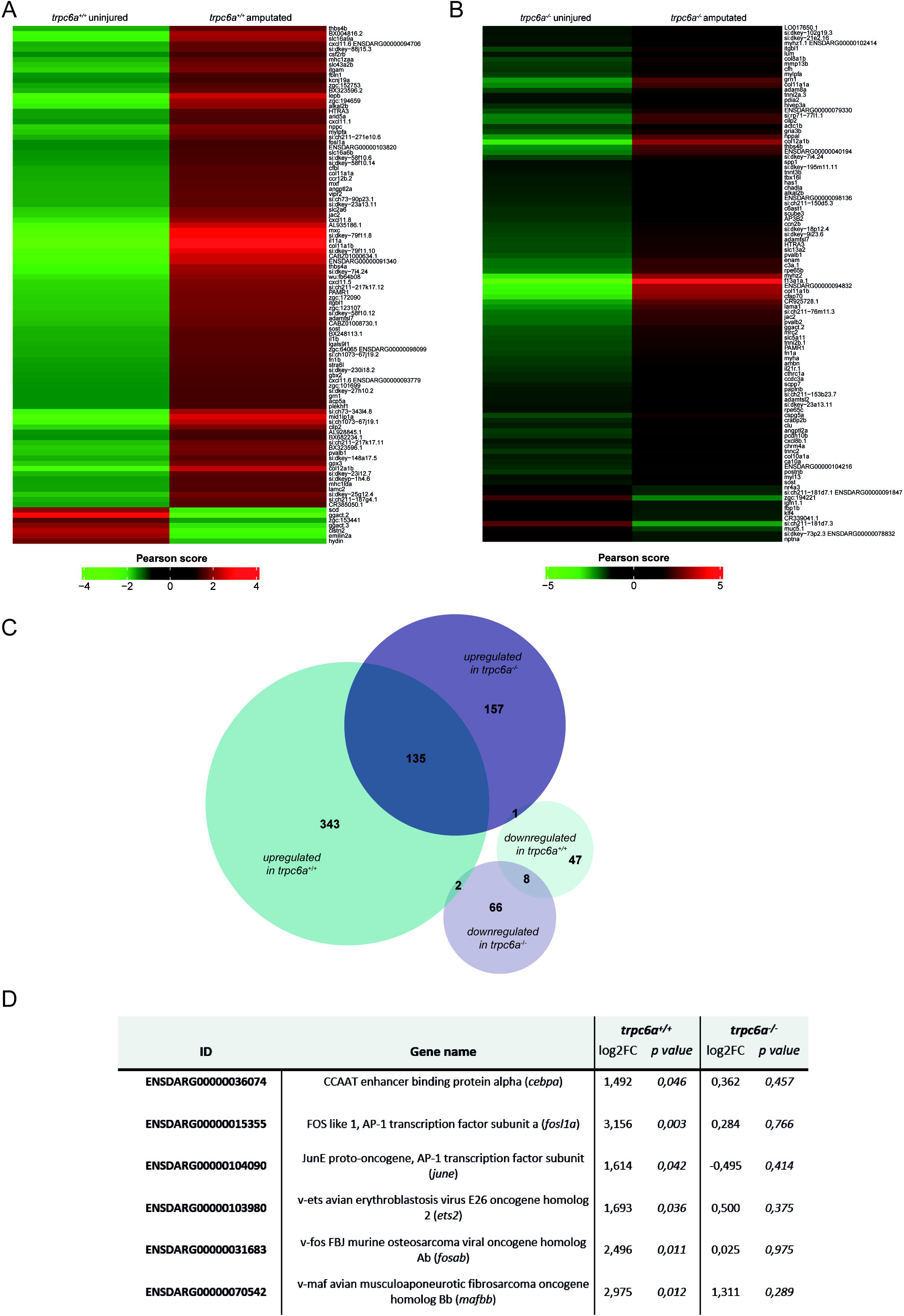
Loss of Trpc6a results in misregulated gene expression during regeneration. **A-B**. Heatmaps showing the 100 most differentially regulated genes between uninjured and 7dpa hearts *of trpc6a*^*+/+*^ (**A**) and *trpc6a*^*-/-*^ (**B**). Color key represents the relative distance between groups and was calculated using Pearson’s correlation coefficient. **C**. Venn diagram representing the number of genes significantly up or down regulated after injury in *trpc6a*^*+/+*^ and *trpc6a*^*-/-*^ hearts. **D**. Table of transcription factors which are significantly upregulated in *trpc6a*^*+/+*^ hearts but not in *trpc6a*^*-/-*^ hearts following injury (compared to their respective sham controls).

### Trpc6a regulates the stretch induced expression of AP1 transcription factor components

Previous data indicates that pathologically stretching cardiomyocytes *in vivo* activates TRPC6, which in turn induces downstream gene expression^11^. Based on this, we assessed whether Trpc6a regulated the expression of the AP1 transcription factor components *fosl1a* and *fosab* in response to mechanical stretch. To achieve this, we dissociated and isolated cardiomyocytes from either *trpc6*^*-/-*^ or *trpc6a*^*+/+*^ hearts and subjected them to 24 hours of cyclic stretch (n=5/group) (Fig.4.A). Following the completion of this protocol, we harvested the cardiomyocytes and performed RT qPCR for both *fosl1a* and *fosab* (Fig.4.A). In this manner, we observed an approximately 5-fold increase in expression of both *fosl1a* and *fosab* in stretched *trpc6a*^*+/+*^ cardiomyocytes when compared to *trpc6*^*-/-*^ cardiomyocytes (Fig.4.B,C). These data indicate that loss of Trpc6a in cardiomyocytes results in a failure to upregulate the expression of the AP1 transcription factor components *fosl1a* and *fosab* in response to mechanical stretch.

**Figure 4.**
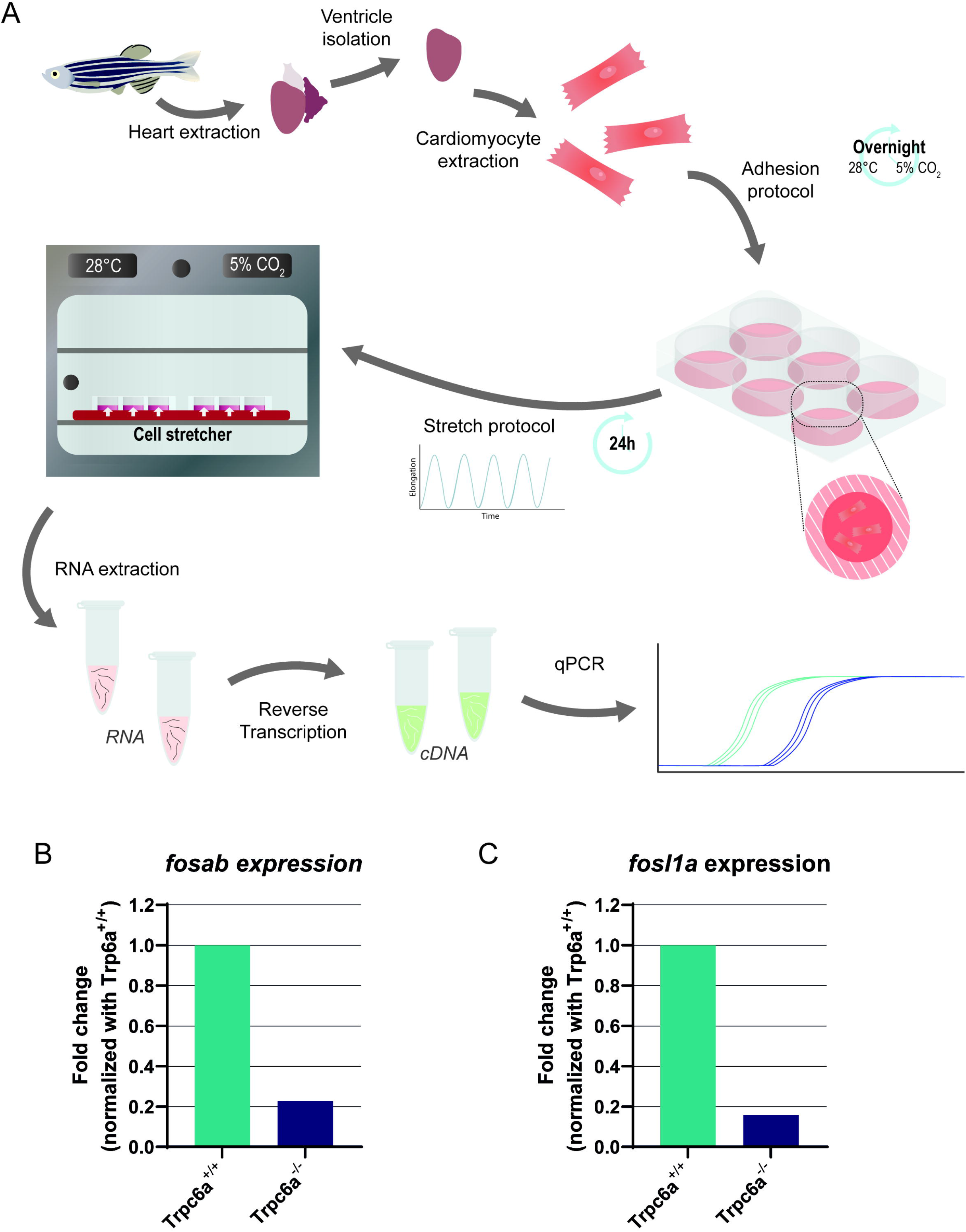
Trpc6a regulates the stretch induced expression of AP1 transcription factor components. **A**. Schematic representation of the experimental design. Cardiomyocytes were isolated from extracted hearts and plated onto Collagen I-coated plates. A static tension protocol was applied overnight to promote cell adhesion and to concentrate cardiomyocytes in the center of the plate. The following day, cell medium was changed and a cyclic stretch protocol was applied for 24h before RNA extraction and RT-qPCR. **B**. Relative expression of *fosab* in *trpc6a*^*+/+*^ and *trpc6a*^*-/-*^ cardiomyocytes subjected to cyclic stretch. **C**. Relative expression of *fosl1a* in *trpc6a*^*+/+*^ and *trpc6a*^*-/-*^ cardiomyocytes subjected to cyclic stretch.

## Discussion

The TRP ion channel TRPC6 is responsible for detecting increased mechanical stretch in cardiomyocytes and activating the CALCINEURIN/NFAT pathway^10^. Under pathophysiological conditions of chronic elevated cardiomyocyte stretch, for example volume overload caused by a cardiac ischemia, this will lead to pathological hypertrophy and ultimately heart failure^10^. Whether TRPC6 could also play a role in detecting increased cardiomyocyte stretch and triggering cardiac regeneration in animal models capable of this feat is currently unknown. The results of this study indicate that Trpc6a is an essential component of the cardiac regenerative response in adult zebrafish. Our data demonstrates that loss of Trpc6a results in a failure to regenerate the heart following cardiac resection. Early regenerative processes such as revascularization appear largely unaffected, however cardiomyocyte proliferation is significantly impeded in the absence of Trp6a signaling leading to the persistence of extensive scarring. Furthermore, comparative transcriptomic analysis of *trpc6a*^*-/-*^ and *trpc6a*^*+/+*^ resected hearts indicates that loss of Trp6a substantially impacts gene expression. In particular components of the AP1 transcription factor complex, which are required for successful cardiac regeneration, are not upregulated in the absence Trp6a. Lastly, our data indicates that Trpc6a regulates the expression of AP1 transcription factor complex components in response to mechanical stretch. Together these results indicate that, in adult zebrafish, increased/chronic cardiomyocyte stretch associated with cardiac injury is sensed by Trpc6a which subsequently activates downstream signaling pathways resulting in the expression of genes, such as AP1 transcription factor complex components, which are involved in driving cardiac regeneration.

The Frank-Starling law was described over a century ago and explains how elevated ventricular preloading, which stretches cardiomyocytes, results in an increased force of contraction in order to maintain circulatory homeostasis^14^. Stretching cardiomyocytes increases the calcium sensitivity of their sarcomeres resulting in enhanced contractility. In situations where ventricular preloading is maintained, there is a further progressive increase in the force of contractility termed the slow force response (SFR) which is driven by elevated, TRPC6 dependent^9^, Ca^2+^ transients^15^. Chronic ventricular loading, for example after myocardial ischemia, results in cardiac remodeling and pathological cardiomyocyte hypertrophy leading, ultimately, to heart failure. The molecular mechanisms which drive pathological cardiac hypertrophy (as opposed to physiological hypertrophy induced by exercise) are largely driven by CALCINEURIN and its downstream effector NFAT^8^. Chronic increases in mechanical load result in elevated intracellular Ca^2+^ which in turn activates CALCINEURIN. CALCINEURIN subsequently dephosphorylates NFAT which translocates into the nucleus and regulates the expression of genes which drive pathological hypertrophy. The ion channel TRPC6 is responsible for detecting increases in cardiomyocyte stretch and generating the sustained Ca^2+^ transients which drive this pathological process^10^. It is apparent then that TRPC6 plays a central role in the cardiac mechanosensitive response to volume overload which results in pathological hypertrophy. Conversely, our data indicates that, in adult zebrafish, it appears that the response to volume overload regulated by Trpc6a results in cardiomyocyte proliferation and ultimately cardiac regeneration. In mice, global KO of *Trpc6* results in increased mortality after myocardial infarction, however this is primarily due to the role Trpc6 plays in cardiac fibroblast avtivation, a process which is essential for early scar formation in order to avoid cardiac rupture^16^. Blood pressure in adult zebrafish is around 50 times lower than mice (2.5mmHg vs 100mmHg)^17^ and as such the formation of a clot is sufficient to avoid excessive blood loss following damage to the myocardium. Because of the role TRPC6 plays in different cell types in mammals, it will be interesting to determine the effect that conditional, cardiomyocyte specific, deletion of *Trpc6* has following myocardial infarction. Conversely, constitutive, cardiomyocyte specific, over-expression of *Trpc6* in mice activates the CALCINEURIN/NFAT pathway resulting in pathological hypertrophy and lethality^10^. However, it would also be of interest to assess what effect *Trpc6* overexpression has at earlier stages of development when cardiomyocytes are still capable of proliferating. Downstream of TRPC6, cardiomyocyte specific overexpression of an active-CALCINEURIN isoform in adult mice is sufficient to trigger pathological hypertrophy and heart failure^18^. Furthermore, similar experiments performed in neonatal mice indicates that active-CALCINEURIN induces a switch from proliferation to hypertrophic growth in cardiomyocytes which are normally hyperplastic at this stage of development^19^. While these data seem at odds with our finding that Trpc6a plays a beneficial role during cardiac regeneration, our transcriptomic data indicates that other Trpc6a mediated mechanisms are also involved. Although much focus has been placed on the TRPC6-CALCINEURIN-NFAT axis, TRPC6 also activates other signalling mechanisms such as AP1 mediated gene transcription^20^. We found that, during cardiac regeneration, there is a significant increase in the expression the AP1 transcription factor components *fosab* and *fosl1a* which does no occur when Trpc6a is absent. This is in-line with previous *in vitro* data indicating that activation of Trpc6 results in increased *c-fos* expression^21^. More recently, the AP1 transcription factor complex has been shown to be a critical regulator of cardiac regeneration in adult zebrafish^12^. In particular, cardiomyocyte specific expression of a dominant-negative *Fos* isoform significantly inhibits cardiac regeneration. This loss of AP1 function results in defective cardiomyocyte sarcomere disassembly and proliferation and also affects their ability to extend protrusions into the site of injury^12^. Interestingly, although the AP1 components *JunB* and *Fosl1* are upregulated in adult zebrafish hearts after injury, the same is not true for adult mice after myocardial infarction^12,22^. Furthermore, overexpression of *JunB* and *Fosl1* in neonatal rat cardiomyocytes is sufficient to induce protrusive behaviour in these cells similar to that observed in zebrafish cardiomyocytes^12^. These data indicate a difference in AP1 signalling between adult zebrafish and mammals which may partly explain the differences we observed following KO of *trpc6a*. Lastly, we have demonstrated that Trpc6a regulates the expression of the AP1 transcription factor components *fosab* and *fosl1a* in response to mechanical stretch, similar to reports in mammalian cardiomyocytes^5,23^. Although care must be taken when extrapolating *in vitro* data, it is likely that this is also the situation which occurs during cardiac regeneration in adult zebrafish. The increase in myocardial stretch caused by volume overload following cardiac injury could activate Trpc6a and induce the expression of AP1 components required for cardiac regeneration. In summary we have identified Trpc6a as a critical regulator of cardiac regeneration which can induce the expression of AP1 components in response to mechanical stretch. Future studies will be required to establish exactly why *trpc6a* induces a regenerative response in adult zebrafish compared to the pathological response in mammals.

## Methods

### Zebrafish line and husbandry

Trpc6a KO G637T (sa23930) zebrafish transgenic line was purchased from ZIRC and maintained under standardized conditions^24^. Experiments were conducted in accordance with local approval and the European Communities council directive 2010/63/EU.

### Larval heart rate and blood flow analysis

3dpf larvae were anaesthetised and mounted in low melt agarose. 30 seconds videos of either cardiac contractions or blood flow were recorded using a Point Grey GRAS-03K2C-C high speed camera. Heart rate and blood flow were analysed using ViewPoint MicroZebraLab software and ImageJ software. T-test statistical analysis was performed using GraphPad Prism.

### Resection

Cardiac resection were performed on 6 to 10-month-old zebrafish as previously described^2^, in accordance with local approval (APAFIS#2021021117336492 v5).

### Immunohistochemistry and histological staining

Immunohistochemistry and histological stainings were performed on 10μm heart sections as previously described^25^. The antibodies used in this manuscript are listed below:

anti-Trpc6 (OST00081W, Osenses)

anti-Trpm (T2780, Sigma)

EdU labelling was performed according to the manufacturer’s instructions (Click-iT EdU Kit C10337, Molecular Probes). Acid Fuchsin-Orange G (AFOG) staining was performed as previously described^26^ and the size of the scar area was calculated using ImageJ software. T-test statistical analyses was performed using GraphPad Prism. Alcaline phosphate staining was performed on whole-mount heart as previously described^25^.

### Imaging

A Zeiss Discovery V20 fluorescence stereomicroscope fitted with a Tucsen FL20 microscope camera was used for histological imaging and either a Zeiss Axio Imager equipped with an Apotome 3 module or a Leica TCS SP-8 confocal microscope were used for imaging immunohistochemistry labelled sections.

### EdU labelling

To label proliferating cells, amputated fish were anesthetized in Tricaine and injected with 50μL of 240μg/mL of EdU solution daily. At 14dpa, fish were euthanized (excess of tricaine), the hearts were collected, and processed for immunohistochemistry. Following imaging, EdU+ cardiomyocytes were counted using IMARIS software. T-test statistical analysis was performed using GraphPad Prism.

### RNA sequencing

Adult fish were anesthetized in Tricaine. For each group, 5 hearts were pooled and RNA was extracted using Trizol/choloform. Genewiz performed the RNAsequencing and bioinformatic analysis of the data using DESeq2. The Wald test was used to generate p-values and log2 fold changes (log2FC). Genes with a p-value < 0.05 were considered differentially expressed. Heatmaps were generated with iDEP 1.0 using the top 100 most variable genes and the distance between groups was calculated using the Pearson’s correlation coefficient.

### Cardiomyocyte isolation and cyclic stretch

For each group, 5 hearts were collected and pooled. Cardiomyocytes were isolated as previously described^27^. Cardiomyocytes were plated on BioFlex® culture plates coated with Collagen I (Flexcell®) and concentrated at the center of the wells using a 12-hour static tension protocol applied with the Flexcell Tension System (FX-6000T, Flexcell®) at 28°C and with 5% CO2. Following this, the culture medium was changed and a cyclic stretch protocol was applied for 24h (Sine, 16% elongation, 0.5Hz).

### Real-time quantitative PCR

RNA was extracted from isolated cardiomyocytes using Trizol/chloroform. cDNA was obtained after reverse transcription using a First strand cDNA synthesis RT-PCR kit(Roche) and quantitative PCR was performed using SYBR Green (Roche) and a LightCycler 480 system (Roche). The primer sequences used are as follow:

*tubulin alpha* Forward: 5′ CGGCCAAGCAACACTACTAGA 3’

*tubulin alpha* Reverse: 5′ AGTTCCCAGCAGGCATTG 3’

*fosl1a* Forward: 5′ AAGGGAACGCAACAAAATGG 3’

*fosl1a* Reverse: 5′ AGCTTCTCCTTTTCCTTCTGG 3’

*fosab* Forward: 5′ GTGCTTTTCGACTTTTGACAGG 3’

*fosab* Reverse: 5′ GTCTGGTTGAGCGGGTAATAC 3’

## Acknowledgements

We would like to acknowledge the imaging facility MRI, member of the France-BioImaging national infrastructure supported by the French National Research Agency (ANR-10-INBS-04, «Investments for the future»)”. We also acknowledge the Zebrafish International Resource Center for providing KO zebrafish lines. The Jopling lab is part of the Laboratory of Excellence Ion Channel Science and Therapeutics supported by a grant from the ANR. Work in the Jopling lab is supported by a grant from the “la Fondation Leducq” and from the ANR (contract ANR-20-CE14-003-02 MetabOx-Heart and ANR-22-CE14-048-02 IONIC).

## References

1. Neves, J. S. et al. Acute Myocardial Response to Stretch: What We (don’t) Know. Front Physiol 6, 408 (2015).

2. Jopling, C. et al. Zebrafish heart regeneration occurs by cardiomyocyte dedifferentiation and proliferation. Nature 464, 606–609 (2010).

3. Porrello, E. R. et al. Transient regenerative potential of the neonatal mouse heart. Science 331, 1078–1080 (2011).

4. Banerjee, I. et al. Cyclic stretch of embryonic cardiomyocytes increases proliferation, growth, and expression while repressing Tgf-beta signaling. Journal of molecular and cellular cardiology 79, 133–44 (2015).

5. Rysä, J., Tokola, H. & Ruskoaho, H. Mechanical stretch induced transcriptomic profiles in cardiac myocytes. Sci Rep 8, 4733 (2018).

6. Yu, J. K. et al. Cardiac regeneration following cryoinjury in the adult zebrafish targets a maturation-specific biomechanical remodeling program. Sci Rep 8, 15661 (2018).

7. Rovira, M., Borràs, D. M., Marques, I. J., Puig, C. & Planas, J. V. Physiological Responses to Swimming-Induced Exercise in the Adult Zebrafish Regenerating Heart. Front Physiol 9, 1362 (2018).

8. Wilkins, B. J. et al. Calcineurin/NFAT coupling participates in pathological, but not physiological, cardiac hypertrophy. Circ Res 94, 110–118 (2004).

9. Yamaguchi, Y., Iribe, G., Nishida, M. & Naruse, K. Role of TRPC3 and TRPC6 channels in the myocardial response to stretch: Linking physiology and pathophysiology. Prog Biophys Mol Biol 130, 264–272 (2017).

10. Kuwahara, K. et al. TRPC6 fulfills a calcineurin signaling circuit during pathologic cardiac remodeling. J Clin Invest 116, 3114–3126 (2006).

11. Seo, K. et al. Combined TRPC3 and TRPC6 blockade by selective small-molecule or genetic deletion inhibits pathological cardiac hypertrophy. Proc Natl Acad Sci U S A 111, 1551–1556 (2014).

12. Beisaw, A. et al. AP-1 Contributes to Chromatin Accessibility to Promote Sarcomere Disassembly and Cardiomyocyte Protrusion During Zebrafish Heart Regeneration. Circ Res 126, 1760–1778 (2020).

13. Ge, R. et al. Critical role of TRPC6 channels in VEGF-mediated angiogenesis. Cancer Lett 283, 43–51 (2009).

14. Patterson, S. W., Piper, H. & Starling, E. H. The regulation of the heart beat. J Physiol 48, 465–513 (1914).

15. Alvarez, B. V., Pérez, N. G., Ennis, I. L., Camilión de Hurtado, M. C. & Cingolani, H. E. Mechanisms underlying the increase in force and Ca(2+) transient that follow stretch of cardiac muscle: a possible explanation of the Anrep effect. Circ Res 85, 716–722 (1999).

16. Davis, J., Burr, A. R., Davis, G. F., Birnbaumer, L. & Molkentin, J. D. A TRPC6-dependent pathway for myofibroblast transdifferentiation and wound healing in vivo. Dev Cell 23, 705–715 (2012).

17. Hu, N., Yost, H. J. & Clark, E. B. Cardiac morphology and blood pressure in the adult zebrafish. Anat Rec 264, 1–12 (2001).

18. Molkentin, J. D. et al. A calcineurin-dependent transcriptional pathway for cardiac hypertrophy. Cell 93, 215–228 (1998).

19. Nguyen, N. U. N. et al. A calcineurin-Hoxb13 axis regulates growth mode of mammalian cardiomyocytes. Nature 582, 271–276 (2020).

20. Thiel, G., Lesch, A., Rubil, S., Backes, T. M. & Rössler, O. G. Regulation of Gene Transcription Following Stimulation of Transient Receptor Potential (TRP) Channels. Int Rev Cell Mol Biol 335, 167–189 (2018).

21. Thiel, G. & Rössler, O. G. Hyperforin activates gene transcription involving transient receptor potential C6 channels. Biochemical Pharmacology 129, 96–107 (2017).

22. van Duijvenboden, K. et al. Conserved NPPB+ Border Zone Switches From MEF2-to AP-1-Driven Gene Program. Circulation 140, 864–879 (2019).

23. Komuro, I. et al. Stretching cardiac myocytes stimulates protooncogene expression. J Biol Chem 265, 3595–3598 (1990).

24. Aleström, P. et al. Zebrafish: Housing and husbandry recommendations. Lab Anim 54, 213–224 (2020).

25. Lai, S.-L. et al. Reciprocal analyses in zebrafish and medaka reveal that harnessing the immune response promotes cardiac regeneration. eLife 6, e25605 (2017).

26. Poss, K. D., Wilson, L. G. & Keating, M. T. Heart Regeneration in Zebrafish. Science 298, 2188–2190 (2002).

27. Sander, V., Suñe, G., Jopling, C., Morera, C. & Izpisua Belmonte, J. C. Isolation and in vitro culture of primary cardiomyocytes from adult zebrafish hearts. Nat Protoc 8, 800–809 (2013).

